# Structure and activity of botulinum neurotoxin X

**DOI:** 10.1101/2023.01.11.523524

**Authors:** Markel Martínez-Carranza, Jana Škerlová, Pyung-Gang Lee, Jie Zhang, Dave Burgin, Mark Elliott, Jules Philippe, Sarah Donald, Fraser Hornby, Linda Henriksson, Geoffrey Masuyer, Matthew Beard, Min Dong, Pål Stenmark

**Affiliations:** Department of Biochemistry and Biophysics, Stockholm University, Stockholm, Sweden; Department of Urology, Boston Children’s Hospital, Boston, MA, USA; Department of Microbiology, Harvard Medical School, Boston, MA, USA; Department of Surgery, Harvard Medical School, Boston, MA, USA; Ipsen Bioinnovation, 102 Park Drive, Abingdon, UK

**Keywords:** Botulinum, neurotoxin, NTNH, BoNT/X, cryo-EM

## Abstract

Botulinum neurotoxins (BoNTs) are the most potent toxins known and are used to treat an increasing number of medical disorders. All BoNTs are naturally co-expressed with a protective partner protein (NTNH) with which they form a 300 kDa complex, to resist acidic and proteolytic attack from the digestive tract. We have previously identified a new botulinum neurotoxin serotype, BoNT/X, that has unique and therapeutically attractive properties. We present the cryo-EM structure of the BoNT/X-NTNH/X complex at 3.1 Å resolution. Unexpectedly, the BoNT/X complex is stable and protease resistant at both neutral and acidic pH and disassembles only in alkaline conditions. Using the stabilizing effect of NTNH, we isolated BoNT/X and showed that it has very low potency both *in vitro* and *in vivo*. Given the high catalytic activity and translocation efficacy of BoNT/X, low activity of the full toxin is likely due to the receptor-binding domain, which presents weak ganglioside binding and exposed hydrophobic surfaces.

## Introduction

Botulinum neurotoxins (BoNTs) are a family of bacterial protein toxins produced by *Clostridium botulinum* and other clostridial bacteria. They are the most potent toxins known and cause flaccid paralysis by targeting motor neurons, translocating their catalytic domain into the cytosol, and cleaving SNARE proteins (soluble N-ethylmaleimide-sensitive factor attachment protein receptors) essential for vesicle – membrane fusion. The affected neurons are rendered unable to release neurotransmitters into the neuromuscular junction, causing muscle paralysis.^1^

BoNTs are 150 kDa proteins and consist of three domains: the 50 kDa catalytically active light chain (LC) and the 100 kDa heavy chain that is further divided into the translocation domain (H_N_) and the receptor-binding domain (H_C_). Structurally, the LC and translocation domain form one closely connected unit, referred to as LH_N_. Canonical BoNTs are classified into seven traditional serotypes, alphabetically named BoNT/A-G.^2^ There are also mosaic toxins composed of domains of different serotypes. A novel serotype named BoNT/X was identified in the genome of *C. botulinum* strain 111, which also contains a BoNT/B gene on a plasmid and was originally isolated from an infant botulism case in Japan.^3–5^ The BoNT/X cluster in *C. botulinum* strain 111 is of the OrfX-type, and encodes an additional OrfX2 protein named OrfX2b.^3^ BoNT/X has emerged as a promising vehicle for intracellular delivery of therapeutic nanobodies, a canvas for engineering substrate specificity, and a potential way to target novel medical conditions.^6–8^ BoNT/X has low sequence similarity to other BoNTs (30% sequence identity) and is not recognized by antisera against other known serotypes. It also exhibits a unique substrate specificity as it cleaves VAMP1/2/3 at a distinct site from other BoNTs, and is the only BoNT capable of cleaving the non-canonical SNAREs VAMP4, VAMP5 and Ykt6. The only structural information available for BoNT/X and its corresponding NTNH to date is the crystal structure of the toxin’s highly active LC.^9^

Recently, several “botulinum neurotoxin-like toxins” have been discovered. These toxins (BoNT/X, BoNT/En and PMP1) are closely related to the established BoNTs but form a distinct evolutionary branch.^3,10,11^ PMP1 is the first toxin in this family to target insects, specifically the malaria vector *Anopheles* mosquitos.^11^ It is not known if BoNT/X or BoNT/En target insects, it should however be noted that these toxins were found in bacterial strains isolated from the human gut or from animal feces, while PMP1 was identified in a mosquitocidal strain isolated from mangrove soil.

The disease botulism is most commonly contracted as a foodborne illness, when the patient ingests poorly conserved food where toxigenic bacteria have grown and produced the neurotoxin. In order to reach the neurons, BoNTs need to resist the acidic environment and proteases present in the gastrointestinal (GI) tract of the host. Several non-toxic neurotoxin-associated proteins (NAPs) encoded in the *bont* gene clusters are known to protect BoNT by forming high molecular weight assemblies.^12^ All known *bont* genes are neighbored in their gene clusters by an *ntnh* NAP gene (nontoxic non-hemagglutinin protein). Together BoNT and NTNH proteins form the 300 kDa minimal progenitor toxin complex (M-PTC). So far the crystal structures of BoNT/A and BoNT/E in complex with their respective NTNH proteins have been solved,^13,14^ revealing a tight complex where NTNH shares the same general fold as BoNT. NTNH is likely a result of a gene duplication event, and has later lost the receptor-binding ability and proteolytic activity.^15^ The BoNT-NTNH complexes are held together by several pH-dependent contacts that release the neurotoxin once the complex leaves the acidic GI tract^16^, and greatly enhance the oral potency of the neurotoxins by 10 to 20-fold compared to BoNT alone.^17^

BoNT/X has successfully been used for delivery of single-domain antibodies into neurons as well as Cas13 and Cas9.^6,8^ The specificity of LC/X has successfully been altered using an innovative phage-assisted evolution method, breaking new ground for the possible uses of the toxins both as therapeutics and as research tools.^7^

Here we have studied the potency of isolated recombinant BoNT/X, determined the structure of the 300 kDa BoNT/X-NTNH/X complex and investigated the complex’s pH-dependent stability. The structure of BoNT/X suggests it is a functional toxin despite its weak activity on the mammalian models tested so far. It provides a template for the design and refinement of BoNT/X-derived biotechnological tools for intracellular delivery of therapeutic molecules or targeted secretion inhibitors.^7,8,18^

## Results and discussion

### Production of active BoNT/X

Full length BoNT/X, and especially the isolated binding domain of BoNT/X (Hc/X), are difficult to express and purify with good yields and purity.^3^ Here, we utilized the stabilizing effect of NTNH on BoNT/X to produce full-length active toxin. We co-expressed BoNT/X and its corresponding NTNH, which allowed us to successfully produce both the recombinant 300 kDa M-PTC complex for structural determination, and the active 150 kDa BoNT/X for *in vitro* and *in vivo* activity studies (Supplementary figure S1). This facilitated a more accurate characterization of this new serotype’s activity and determination of the structure and pH stability of the BoNT/X-NTNH/X complex (M-PTC/X complex).

### Activity of BoNT/X in cultured neurons

First, the ability of BoNT/X to reach and cleave intracellular VAMP was assessed in rat cortical neurons (Figure 1). Proteolytic degradation was observed for VAMP2 and VAMP4, with 10 nM BoNT/X able to cleave 58% of both substrates (Figure 1 and Supplementary figure S2). Although this activity shows BoNT/X is functional on neurons, it appears considerably less potent than other serotypes. For comparison, BoNT/B presented a sub-picomolar EC_50_ when analyzing VAMP2 cleavage in rat spinal cord neurons,^19^ which was similar to BoNT/A cleavage of SNAP25 in rat cortical neurons.^20^ Remarkably, BoNT/X was only marginally more efficient than LH_N_/X, a fragment that lacks the receptor-binding domain, and was able to degrade 42% of intracellular VAMP2. Previous reports evaluating the functionality of LH_N_ showed that it could retain some level of activity on cultured neurons, but was approximately 10^5^-fold less potent than the full-length BoNT/A and B toxins.^19,21^ The activity observed here with LH_N_/X suggests this fragment also has the intrinsic ability to transport LC/X inside neuronal cells, within a similar range to what was observed for LH_N_/B, which presented an EC_50_ of 15 nM for VAMP2 cleavage. This is consistent with previous observations where association of LH_N_/X with the binding domain of BoNT/A formed a chimeric toxin platform for intracellular delivery of cargo proteins.^8^

**Figure 1:**
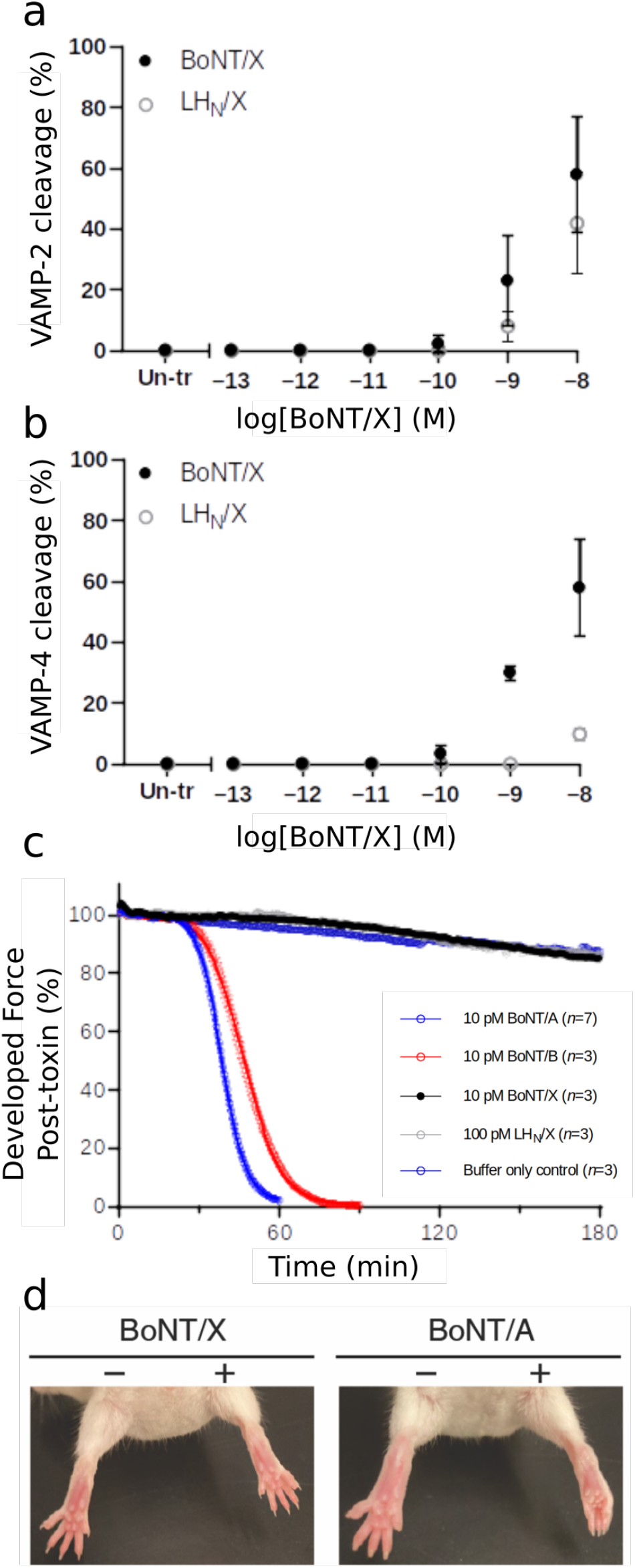
Activity of BoNT/X. Rat cortical neurons were exposed to varying concentrations of recombinant full-length BoNT/X (filled circles) or LH_N_X (open circles). After 24 h of incubation cells were lysed, and lysates analyzed for VAMP2 (**a**) and VAMP4 (**b**) by Western blots (n=3). (**c**) Mouse phrenic nerve hemi-diaphragm (mPNHD) preparations were exposed to 10 pM BoNT/X, BoNT/A and BoNT/B, 100 pM LH_N_/X, and muscle contraction responses to indirect stimulation recorded until contraction was no longer detectable or to 180 min (n=3). (**d**) BoNT/A (6 pg) induced flaccid paralysis when injected into the right gastrocnemius muscle of mice in DAS assays, whereas injection of BoNT/X (1 μg) did not induce muscle paralysis.

### Activity of BoNT/X in animal models

In the original characterization of BoNT/X,^3^ no active holotoxin was produced due to safety considerations. However, an initial assessment of its toxicity was performed by synthesizing a limited amount of complete toxin using sortase-mediated ligation of the isolated functional LH_N_ and H_C_ domains. Although H_C_/X was observed to be relatively unstable on its own, which lead to a low yield of ligated toxin, the product from the sortase reaction caused local flaccid paralysis in a mice Digit Abduction Score (DAS) assays, when administered at high concentrations.

Here we analyzed the activity of full-length BoNT/X holotoxin (not sortase ligated) utilizing both the mouse phrenic nerve hemi-diaphragm assay and the DAS assay. The mouse phrenic nerve hemi-diaphragm assay provides a useful *ex vivo* assessment on the effects of BoNTs by measuring a decrease in the contraction amplitude of the indirectly stimulated muscle.^22^ BoNT/X was directly compared to BoNT/A and BoNT/B. As expected, the two controls, 10 pM BoNT/A and BoNT/B produced a time-dependent paralysis of the hemi-diaphragm muscle (t_50_=39±0.8 mins, and 47±1.5 mins (n=3), respectively). In contrast, an equivalent dose of BoNT/X (10 pM, n=3) had no effect on diaphragm muscle contraction, nor did LH_N_X at a higher dose (100 pM) (Figure 1c), thus suggesting that the low potency observed *in vitro* is too weak to translate into animal toxicity.

Next, BoNT/X holotoxin was evaluated in the DAS assay, which is commonly used to measure the paralytic effects of BoNTs on muscles *in vivo*.^23^ Injections of 1 or 2 μg of BoNT/X in mice did not cause any paralysis (Figure 1d and supplementary table 1 and 2). This shows the lack of toxicity of BoNT/X, since typical doses for BoNT/A and /B are normally within the pM range to observe local paralysis of the leg muscle. In addition, the mice body weight was not affected by injection of 1 μg of BoNT/X (Supplementary table 3), suggesting that the toxin does not cause significant systemic effect at this dose. Previously, it was reported that 0.5 μg of sortase-ligated toxin resulted in discernable paralysis in the DAS assay, although intraperitoneal injection of 1 μg did not cause any observable effect.^3^ The discrepancy in the DAS assay might be due to some inherent instability of the recombinant toxin compared to the synthetic ligated protein, or it is possible that the considerable amount of free LH_N_/X leftover in the ligation sample contributed to the original observed muscle paralysis.

### Cryo-EM structure of BoNT/X in complex with NTNH (M-PTC/X)

We have determined the cryo-EM structure of the M-PTC/X complex comprising the BoNT/X holotoxin and its non-toxic interaction partner NTNH/X at a nominal resolution of 3.1 Å (using the gold-standard FSC criterion of 0.143), obtained from 432,063 particles selected from a total of 5,408 recorded movies from two datasets in similar data acquisition settings, described in the methods section (see Supplementary Figure S3 and Supplementary table 4). The overall structure of the BoNT/X-NTNH/X complex is shown in Figure 2a-b. Figure 2c shows an assessment of the local resolution throughout the entire map, demonstrating that the core of the complex has a resolution better than 3.1 Å. Figures 2d-f provide visual examples of the map quality.

**Figure 2.**
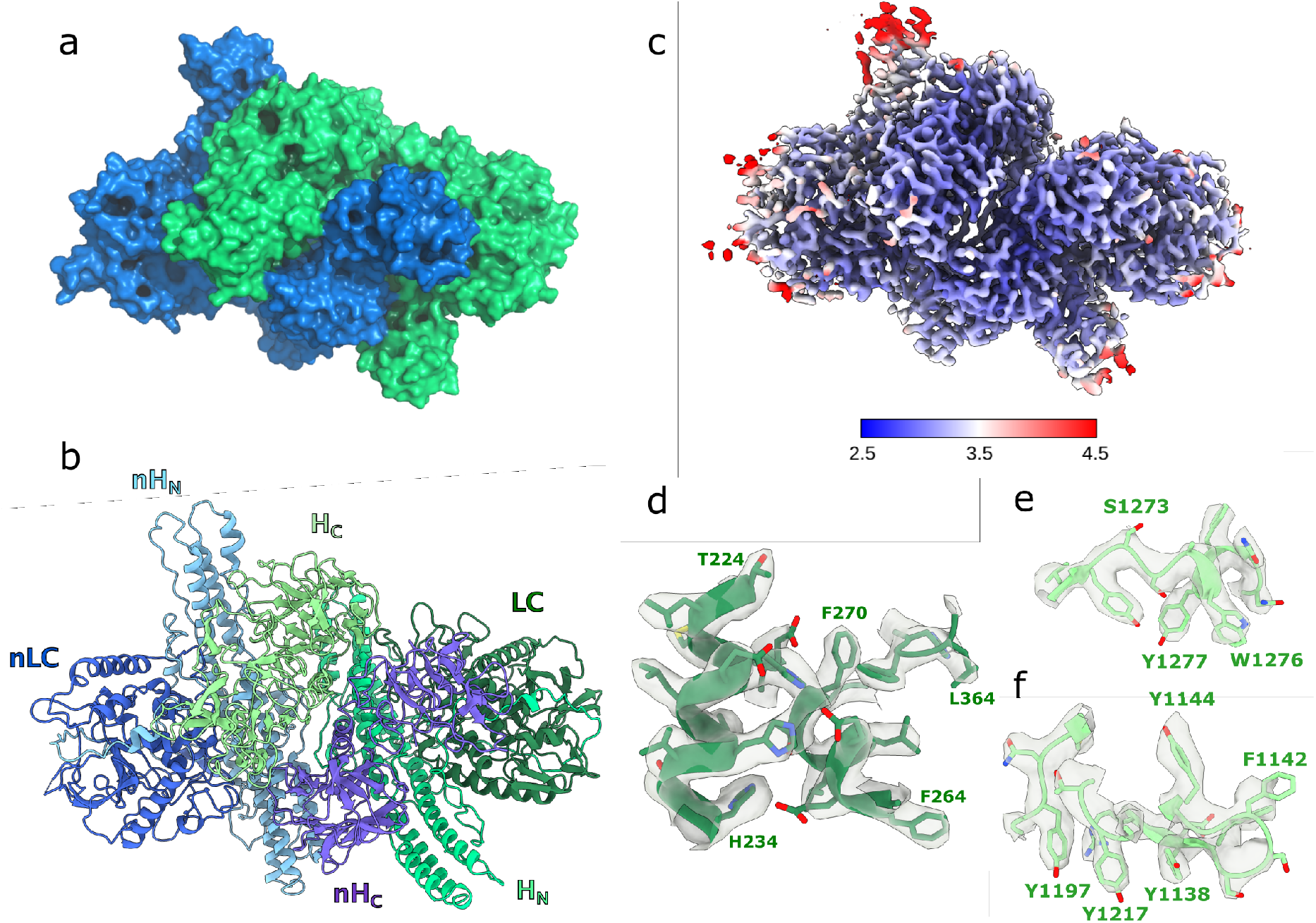
(**a**) Surface view of the BoNT/X (green) – NTNH (blue) complex. (**b**) Cartoon view of the different domains in each protein. BoNT/X domains are colored in green shades, NTNH/X domains are colored in blue shades. (**c**) Local resolution estimates (Å) for the cryo-EM map of the M-PTC/X. (**d**) cryo-EM map around the LC active site. (**e**) cryo-EM map around the SxWY ganglioside-binding motif. (**f**) cryo-EM map around the H_C_ patch rich in aromatic residues.

BoNT/X and NTNH/X share a similar fold and their overall domain arrangement and molecular architecture is similar to other toxin serotypes (Figure 2a-b and Supplementary Figure S4). Similar to the known M-PTC structures of the A and E serotypes, BoNT/X and NTNH/X form an interlocked complex that buries a large solvent-accessible area of 4,778 Å^2^, which is significantly larger compared to BoNT/A-NTNH/A (3,664 Å^2^) and BoNT/E-NTNH/E (3,459 Å^2^). The main domains that are involved in this interaction are the receptor-binding domain in BoNT/X (H_C_/X) and the analogous domain in NTNH/X (nH_C_/X), together with the translocation domain and its analogue (H_N_/X and nH_N_/X). LC and nLC are situated in both extremes of the complex, facing opposite directions from the interface and are mostly solvent-exposed. The critical disulfide bond between C423 and C467 that holds the light and heavy chains of the toxin together after nicking by host or bacterial proteases is discernible in the map, in the same position as in the structures of BoNT/A, /B, and /E.^13,14,24^ It has been shown that C461 can also form a disulfide bond with C423, with the toxin retaining activity in C467S mutants.^3^ Our cryo-EM map indicates that C423-C467 is the most likely natural disulfide bridge. C461 is not visible in the map, as it is located in a loop which is either nicked or disordered. In our structural studies we used a catalytically inactive double mutant construct of BoNT/X described in the methods section. The mutated residues are not involved in the coordination of the Zn^2+^ atom, and the map in the area between residues E226, H227 and H231 indicates that the metal likely is present in the active site. At this resolution we decided not to include it in the model since we previously have determined the 1.35 Å crystal structure of the catalytic domain of BoNT/X, describing the metal coordination in details.^9^

NTNH/X lacks the nLoop conserved in the nLC of NTNH/A1, B, C, D, and G, which facilitates the assembly of the L-PTC with the HA proteins (Supplementary Figure S5).^25–27^ This correlates with the absence of HA proteins in the BoNT/X gene cluster, which is of the *orfX*-type^3^. However, NTNHX has an extended loop with exposed hydrophobic residues, structurally close to the NTNHA nLoop. Although not conserved in other *orfX*-type NTNTH proteins, its position suggests a potential interaction site with other proteins of the BoNT/X cluster.

The translocation domain consists of two central coiled-coil helices of approximately 105 Å flanked by smaller helices and linked with LC through a region known as the ‘belt’. The belt of BoNT/X wraps around LC (Figure 3a), likely acting as a molecular chaperone, as has been shown for BoNT/A.^28^ It occupies part of a groove on the LC surface which is also associated with substrate-binding.^29^ The belt adapts remarkably to the LC surface, as its presence in the full-length toxin does not affect the local structure of LC with interacting regions presenting similar conformations to those seen in the free domain structure (Figure 3c). As in other BoNTs, the extended interaction site of the belt region with LC/X is likely to also mimic the binding of LC to its substrate proteins (VAMP1,2,3,4,5 and Ykt6) which are expected to involve multiple exosites away from the catalytic center (Figure 3a).^9,29^ Figure 3b shows the overlap between a VAMP-like inhibitor bound to LC/F and the H_N_/X belt region.^30^

**Figure 3.**
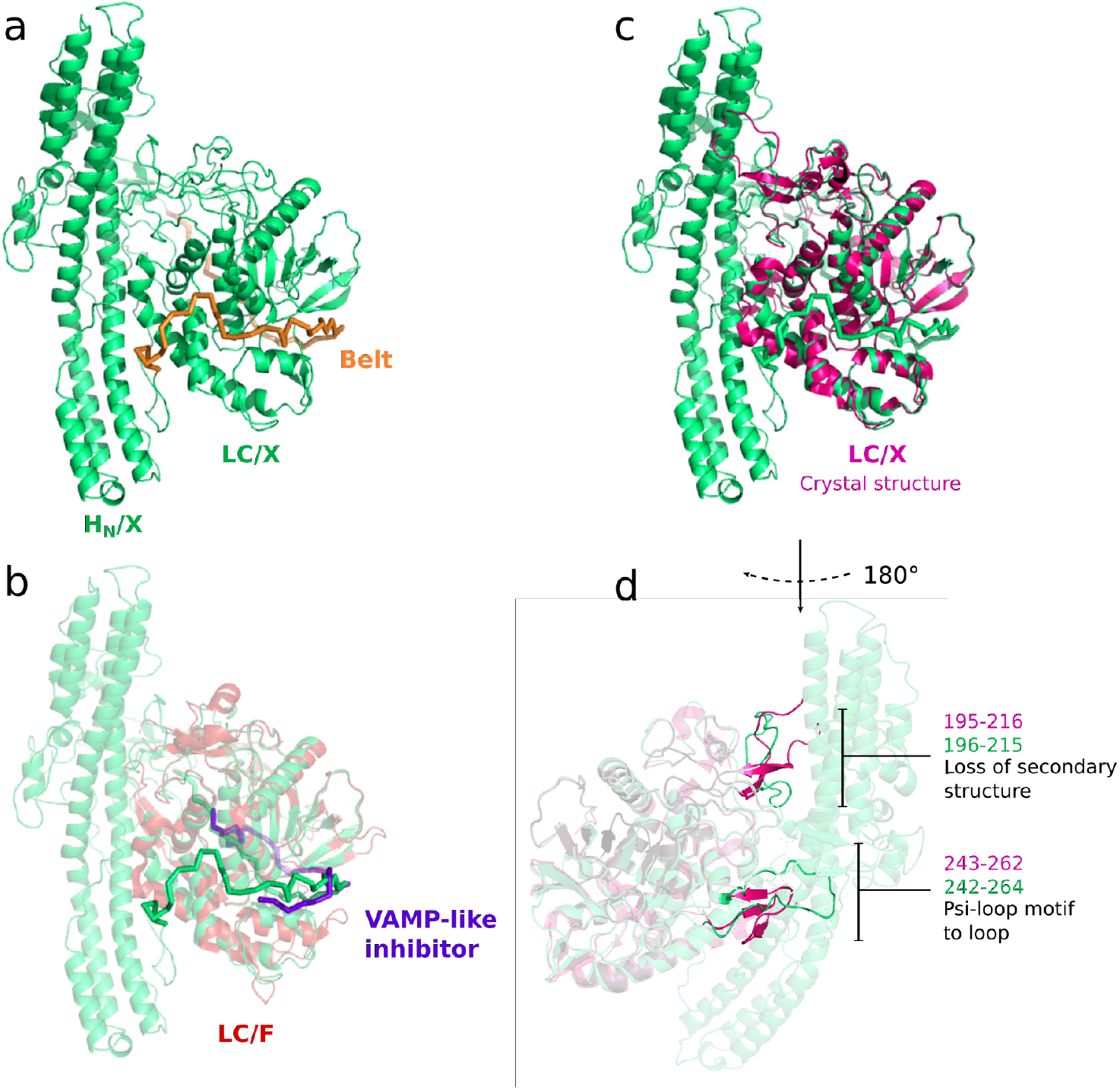
(**a**) H_N_ and LC domains from the BoNT/X-NTNH/X cryo-EM structure (green). The belt region of H_N_ surrounding the LC is highlighted in orange. (b) crystal structure of LC/F (PDB ID 3FIE, colored in red) bound to a VAMP substrate-like inhibitor (purple) superimposed to H_N_-LC/X (green). (**c, d**) crystal structure of LC/X (PDB ID 6F47, colored in magenta) superimposed to the cryo-EM structure of H_N_-LC/X (green). Noteworthy differences in secondary structure are highlighted in panel (**d**).

The translocation domain shares an extensive interaction with LC beyond the belt, and presence of the central H_N_ has a significant impact on the LC structure when compared to the free domain, as shown in Figure 3d. On one side, the H_N_ C-terminal linker (875-878) interacts with LC to slightly rearrange loop 272-278 *via* hydrophobic interactions between P877/F878 and I273. Further, what appears as an extended loop (195-215) stabilized by two β-strands in the free LC, is seen with a much more compact conformation in the complex, with loss of the secondary structure as the I207-V208 residues are pushed by hydrophobic interaction with Y895 and Y951 on the binding domain (H_CN_). These interactions between LC and H_CN_ are interesting as they might also help to stabilize the binding domain in a similar position in the free holotoxin. Furthermore, a significant structural change was also observed in the vicinity of the catalytic site where loop 243-263 takes on a long and flexible conformation extending away from the catalytic pocket, so that T255 interacts with H_N_. In the free LC/X structure, residues 243-263 take on a more complex psi-loop motif in close proximity to the catalytic pocket entrance. The plasticity of this loop may be important for substrate recognition.

### Analysis of the complex’s pH-dependent stability

In order to release the neurotoxin upon leaving the gastrointestinal tract, the M-PTC needs to disassemble in a pH-dependent manner. The large interface in M-PTC/X contains several interaction patches that are displayed in Figure 4a. The interface patches in BoNT/X (theoretical isoelectric point of 5.6) are predominantly positively charged and those in NTNH/X (theoretical isoelectric point of 4.6) are mostly negatively charged. The M-PTC/X is therefore stabilized by a number of electrostatic interactions between BoNT/X and NTNH/X shown in Figure 4b. A cluster of solvent-accessible acidic and basic residues can be found on the interface between H_CN_/X and nH_N_/X, namely D998, K1000, K1046, E1048, E1049, K1050, D1051, K1013, and D1018 in BoNT/X; and K636, E787, E789, and E793 in NTNH/X. The location of this pH-sensing cluster corresponds to a similar cluster in BoNT/A-NTNH/A, which has been demonstrated to drive the pH-mediated disassembly of the M-PTC/A.^13^ The charge of these residues will depend on the environmental pH, which was pH 5.5 in the buffer utilized for the cryo-EM experiments. It is not trivial to predict the pH at which the protonation state of the residues in this pH-sensing cluster will change, since the p*K*_a_ of the amino acids strongly depends on their local chemical environment.^31^

**Figure 4.**
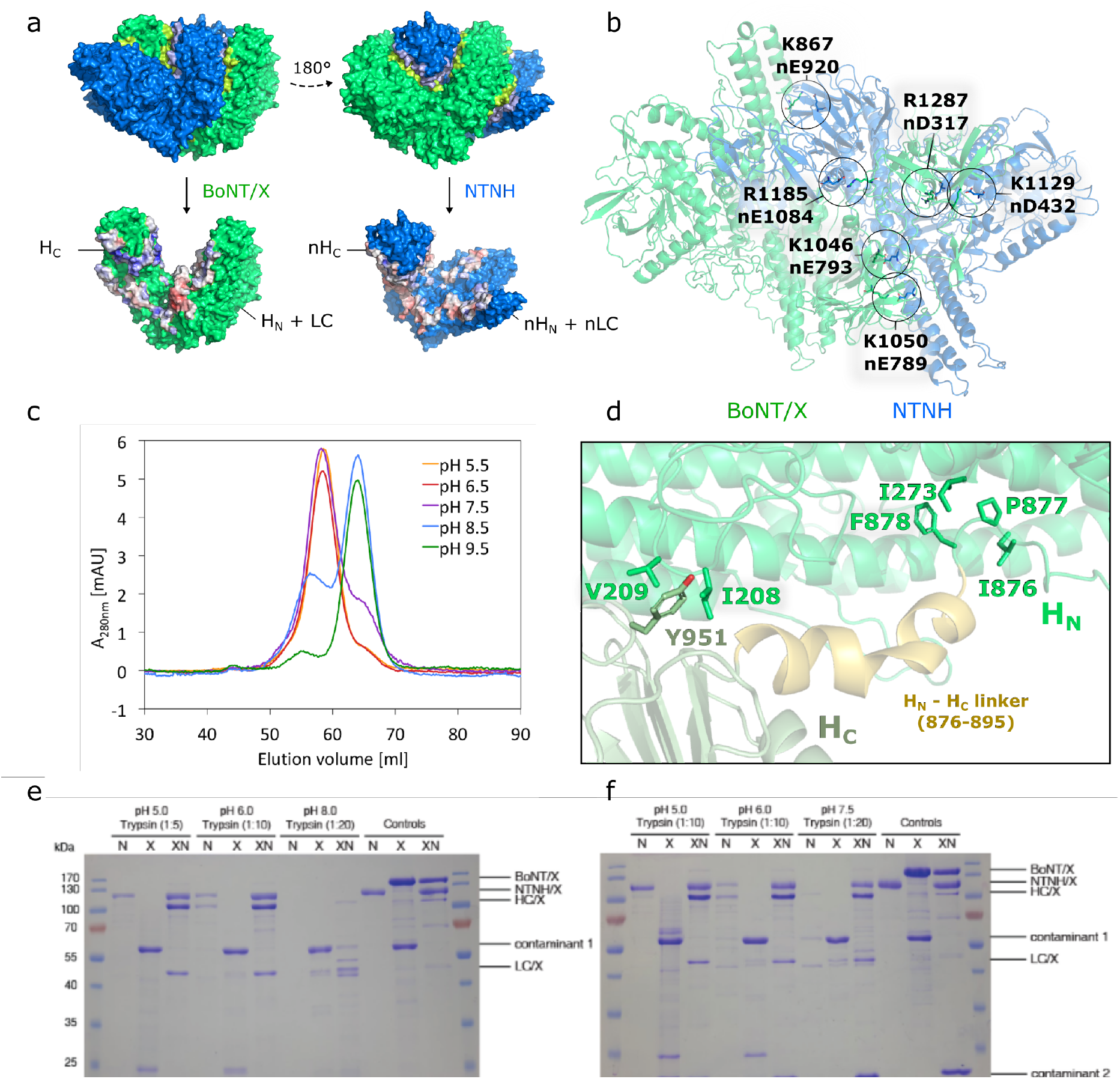
(**a**) BoNT/X-NTNH/X interface; BoNT/X is shown in green and NTNH/X in blue. Top: interfacing residues are highlighted in different shades of green and blue. Bottom: interface patches are colored according to electrostatic potential (negative electrostatic potential is colored in shades of red, and positive in shades of blue). (**b**) Electrostatic interactions across the BoNT/X-NTNH/X interface. (**c**) Size-exclusion chromatograms from the M-PTC/X pH stability experiments. The A_280nm_ response for the chromatograms at pH 5.5 and 6.5 was divided by the factor of 2 to normalize it to the other chromatograms, where the signal was lower. (**d**) Flexible linker between H_N_ and H_C_ followed by a small helical region leading to the H_C_ (residues 876-895) (**e** and **f**) Limited proteolysis assay for examining the stability of the BoNT/X-NTNH/X complex under different pH conditions. BoNT/X (marked as X), NTNH/X (marked as N), and BoNT/X-NTNH/X complex (marked as XN) were incubated with trypsin under the indicated pH conditions and then analyzed by SDS-PAGE and visualized by Coomassie blue staining. BoNT/X alone and NTNH/X alone were degraded under all pH conditions. BoNT/X-NTNH/X is largely resistant to trypsin under pH 5.0, 6.0, and 7.5, but not under pH 8.0. Trypsin treatment is able to cleave the linker between LC and HC of BoNT/X within the BoNT/X-NTNH/X complex, thus BoNT/X showed as two separate bands (LC/X and HC/X) after the disulfide bond connecting the LC and HC is reduced. One contaminant is present in purified BoNT/X and another is present in purified BoNT/X-NTNH/X complex.

In order to determine the pH at which the complex disassembles, we analyzed size-exclusion chromatography profiles of BoNT/X-NTNH/X at pH 5.5, 6.5, 7.5, 8.5, and 9.5. The chromatograms are shown in Figure 4c. The two peaks in the chromatograms correspond to the BoNT/X-NTNH/X complex and to a mixture of free BoNT/X and free NTNH/X, respectively. The complex is stable at pH 5.5, 6.5 and 7.5. At pH 8.5, the equilibrium is shifted towards free BoNT/X and NTNH/X, and at pH 9.5 the complex is almost totally disassembled. To further evaluate the stability and function of the BoNT/X-NTNH/X complex, we then carried out limited proteolysis studies with trypsin treatment under a range of pH conditions (pH 5.0, 6.0, 7.5, and 8.0). As shown in Figure 4e-f, we found that trypsin easily degraded BoNT/X and NTNH/X under all pH conditions, whereas the BoNT/X-NTNH/X complex is largely resistant to trypsin under pH 5.0, 6.0, and 7.5, demonstrating that forming the complex protects BoNT/X from proteases under these pH conditions. After trypsin treatment, BoNT/X within the complex is separated into two fragments of ~100 and ~50 kDa on SDS-PAGE gels (Figure 4e-f): this is likely the HC and LC of BoNT/X as trypsin can still cleave the linker region between the HC and LC.^32^ This protection from trypsin was not observed under pH 8.0 for the complex, suggesting that the complex is not stable under basic conditions, which is consistent with our analysis by size-exclusion chromatography.

Dissociation of the BoNT/X-NTNH/X complex occurs at a significantly more basic pH than BoNT/A-NTNH/A, which is stable at pH 6.0 but disassembled at pH 7.5.^13^ These results, together with its relatively low toxicity in mice, make it tempting to speculate that BoNT/X might target an organism with a different gut environment. Insects for instance have a more alkaline digestive system, and anopheles mosquitoes are known to be the target of the BoNT-like PMP1.^11^ However, BoNT/X was discovered in a clostridial strain associated to a clinical case of human botulism.

### Characterization of the receptor-binding domain of BoNT/X

Although BoNT/X was shown to induce flaccid paralysis in mice at high concentration, its neuronal receptors remain elusive^3^. The M-PTC/X structure presented in this study provides the first insights of the toxin’s receptor-binding domain (H_C_/X). Overall, H_C_/X presents the same architecture observed across all clostridial neurotoxins, which include the lectin-like fold of the N-terminal subdomain (H_CN_/X), which is essential for toxicity,^33^ and the C-terminal subdomain (H_CC_/X). This β-trefoil fold is the main element responsible for neuronal recognition.

The binding domain position has been observed to vary in the different holotoxins, from a linear configuration of the three functional domains in BoNT/A and B to a more compact, closed, structure in BoNT/E where the H_C_ interacts with both the H_N_ and the LC.^34^ This unique position of H_C_/E was suggested to promote the faster translocation rate of BoNT/E.^35^ Remarkably, the position of the H_C_ within the M-PTC is consistent across all serotypes, where it is stabilized by its interaction with NTNH into an intermediate configuration (supplementary figure S4).^13,14^ Mobility of the binding domain is supported by a flexible linker between H_N_ and H_C_, which involves a short helical component (Figure 4d). Although there is no particular sequence conservation for that region, BoNT/X in the M-PTC structure also presents a flexible linker followed by a small helical conformation leading to H_C_ (residues 876-895). Interestingly, this segment is located between the two LC and H_C_ interaction sites that were mentioned previously, and that mainly involve hydrophobic interactions (Figure 4d). This suggests that the domain organization of the free toxin may be similar to the one observed in the M-PTC.

Canonical BoNTs (BoNT/A-BoNT/G) and the tetanus neurotoxin all have gangliosides as receptors or co-receptors. The toxins recognize these cell surface carbohydrates through a shallow pocket centered around the conserved SxWY ganglioside binding motif on H_CC_,^36^ which is also present in the primary sequence of BoNT/X^3^. Superposition of H_C_/X from our structure with ganglioside-bound H_C_/A (PDB ID 5TPC, rmsd of 1.711 Å over 388 Cα pairs, Figure 5a) reveals that the SxWY motif is oriented similarly to that in H_C_/A, suggesting it could accommodate a carbohydrate moiety. However, the variation in ganglioside specificity and affinity observed in BoNTs is mainly driven by non-conserved residues surrounding the common binding motif,^37^ making it difficult to predict a preferred ganglioside receptor for BoNT/X.

**Figure 5.**
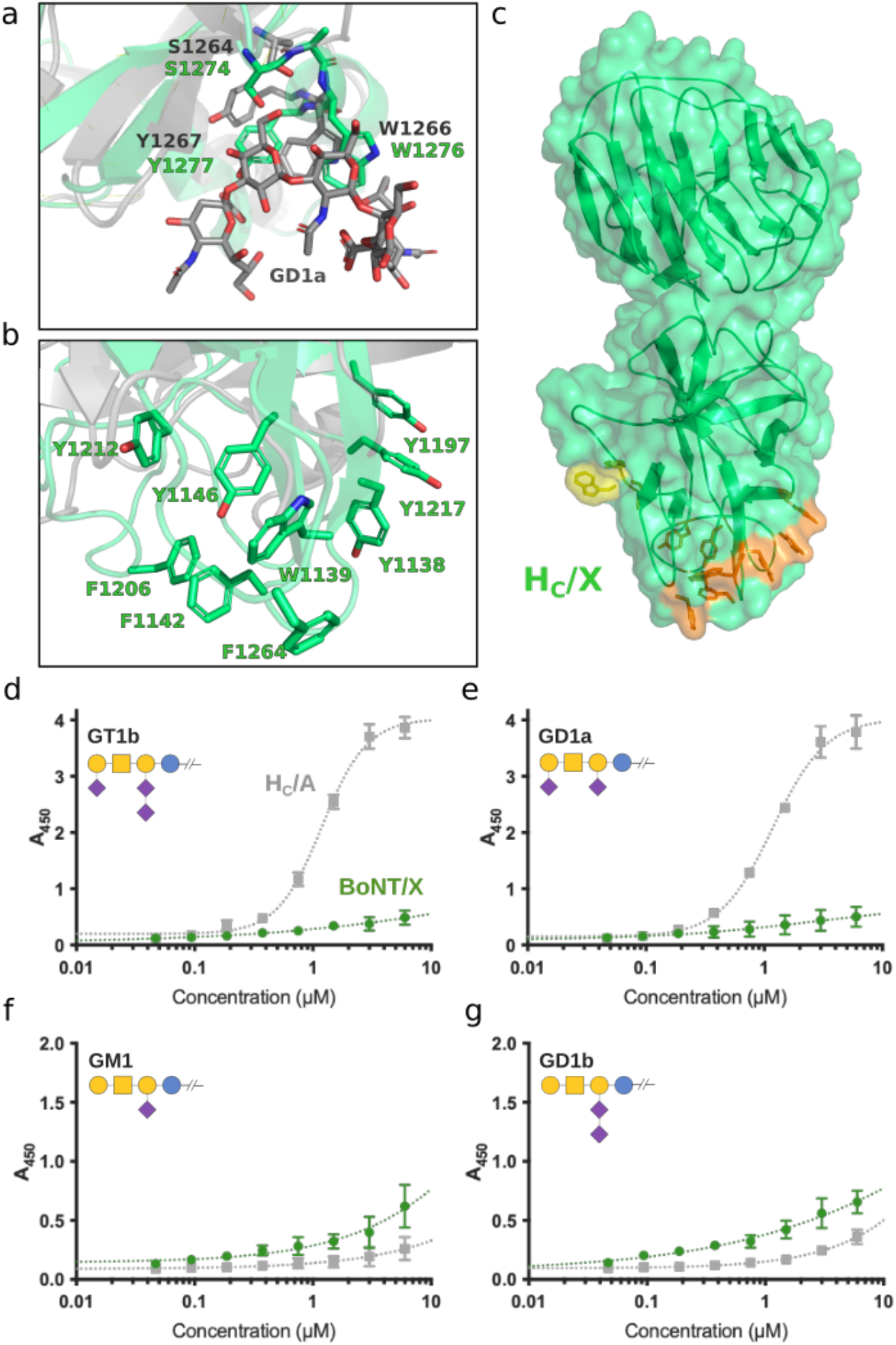
(**a**) Alignment of H_C_/X (green) and GD1a-bound H_C_/A (in grey, PDB ID 5TPC), showing the conserved ganglioside-binding SxWY motif residues in sticks and (**b**) the same alignment showing the exposed region rich in aromatic residues. Location of the ganglioside-binding motif (yellow) and the hydrophobic patch (orange) is shown in the context of H_c_/X in panel (**c**). Binding of BoNT/X (green), and H_C_/A (grey) to GT1b (**d**), GD1a (**e**), GM1 (**f**), and GD1b (**g**), n=3. Glycobloc schematic representation is shown for each ganglioside.^38^

Binding of BoNT/X to gangliosides GT1b, GD1b, GD1a and GM1 was analyzed (Figure 5d-g) using a plate-based assay as described previously.^39,40^ BoNT/X was used rather than H_C_/X because of the reported instability of this domain when expressed individually. Despite presenting the consensus SxWY motif, BoNT/X showed very weak binding to all assayed molecules, which are the most abundant mammalian neuronal gangliosides.^41^ As expected, the BoNT/A1 control presented strong binding to GT1b and GD1a, but not to GD1b and GM1.^42^ Interestingly BoNT/X presented a consistently higher affinity than BoNT/A1 for gangliosides that lack the Sia5 moiety (GD1b and GM1) (Figure 5). Although these results are not entirely conclusive, they hint at a natural carbohydrate-binding capacity for BoNT/X. Ganglioside recognition is considered an essential first step in BoNT toxicity as it provides abundant anchorage on neuronal cell surface where the toxin can then attach to their high-affinity protein receptors for uptake. ^36^ The weak binding of BoNT/X might help explain the low toxicity observed so far in mammals.

BoNT serotypes A,^43–45^ D,^46^, E^47^, and potentially F^39,48^ are known to bind to synaptic vesicle glycoprotein 2 (SV2). The only structural details available to date of the interaction with SV2 are with type A, through the crystal structures of its binding domain (H_C_/A) in complex with the luminal loop 4 of SV2C.^45,49–51^ The H_C_/X backbone superposes well overall with the glycosylated SV2C-bound H_C_/A (PDB ID 5JLV), presenting an rmsd of 1.915 Å over 388 Cα pairs, differing mostly in the H_CN_ region. In H_C_/A, SV2 recognition occurs through a complementary beta-sheet between the SV2C and a beta-hairpin loop of H_C_. Notably, the corresponding site in BoNT/X shows no sequence conservation and no analogous secondary structure. In addition, residues F953 and H1064 of H_C_/A that form stacking interactions with the N559-glycan of glycosylated SV2C, which was shown to be essential for neuronal recognition, are also not conserved in H_C_/X.

Structurally, H_C_/X is more similar to H_C_/D, H_C_/E and H_C_/F than H_C_/A, however, there is no obvious conservation of a potential binding site for SV2. Rmsd values for these superpositions are summarized in supplementary table 5.

The other principal BoNT protein receptors identified so far are synaptotagmins I and II for BoNT/B, DC, and G.^52–57^ BoNT/B and DC bind to synaptotagmin with varying affinity and at two distinct sites located perpendicularly to each other.^58–60^ H_C_/X would need to undergo significant conformational changes in order to accommodate synaptotagmin at the equivalent sites, since loop 1139-1144 overlaps with the H_C_/B synaptotagmin binding site, and loops 1201-1209 and 1247-1252 occupy the H_C_/DC synaptotagmin binding site. The structures of free and synaptotagmin-bound H_C_/B only show minor structural differences (agreeing with an rmsd of 0.566 Å over 441 Cα pairs, PDB IDs 2NM1 and 1Z0H),^58,59,61^ suggesting that a significant conformational change upon receptor binding is unlikely in H_C_/X.

Remarkably, H_C_/X has a pronounced patch of exposed aromatic and hydrophobic residues, which is thermodynamically very unfavorable and uncommon in soluble proteins unless it has an important function. This hydrophobic patch on the H_C_/X loops, including residues Y1138, W1139, F1142, Y1146, Y1197, F1206, Y1212, Y1217, and F1264 (shown in Figure 5b,c), may be involved in neuronal recognition via interactions with a protein or glycosphingolipid receptor, or with the phospholipid-bilayer of the neuronal membrane itself. This hydrophobic patch in BoNT/X is located at the same position in the binding domain as where BoNT/B binds its protein receptor.^58,59,62^ Interestingly, PMP1 has a pronounced hydrophobic patch with similar properties, located at a similar position in the binding domain as BoNT/X, even though the sequence conservation is low.^11^

Hydrophobic interactions directly with the membrane have recently been described as complementing the receptor binding capacity of BoNT/B, /G, and /DC.^1,42,58,59,63,64^ However, the hydrophobic patch on the binding domains of PMP1 and BoNT/X are structurally distinct from the single, extended, hydrophobic loop that contributes to anchoring these toxin serotypes to the membrane.

### Concluding remarks

In this study we solved the cryo-EM structure of the 300 kDa BoNT/X-NTNH/X complex, which is the first M-PTC structure solved by this method. We provide a template for the structural analysis of these complexes using single particle cryo-EM, and demonstrate its usefulness for future studies of other BoNT-NTNH systems. We were able to reach similar resolution to the X-ray crystallography structures determined for BoNT/A-NTNH/A and BoNT/E-NTNH/E, without the significant challenge of obtaining crystals of sufficient quality for these large complexes or the use of nanobodies to facilitate crystallization.^13,14^ The BoNT/X-NTNH/X complex structure also provides structural information on the receptor-binding domain of BoNT/X, which revealed an unusual array of surface-accessible hydrophobic patches despite presenting the same overall fold as other BoNTs. Although H_C_/X presents a conserved carbohydrate-binding motif, it did not bind significantly to any of the most abundant human gangliosides. The structure hence hints at a unique mechanism of cell recognition.

Stability of the M-PTC is known to be pH-dependent to promote oral toxicity and release of the toxin once it has crossed the intestinal barrier. Assessment by size-exclusion chromatography and protease assays showed that the complex remains stable up to pH 7.5, which is an unusual property contrasting with BoNT/A-NTNH/A that dissociates at a lower pH. The structure revealed patches of acidic and basic residues at the interface between H_C_/X and nH_N_/X that are likely to substantially contribute to the pH-sensing mechanism of the complex disassembly. NTNH certainly provides stability for BoNT/X and thus the M-PTC could be a valuable component in the production and formulations of BoNT/X-derived biologicals.

BoNT/X presented very weak activity on cultured neurons and *in vivo*, even more so than in preliminary reports^3^. The LC of BoNT/X was shown to have a particularly high catalytic activity,^9^ and here the LH_N_/X fragment was shown to retain a functional level of toxicity on neuronal cells which was comparable to the same fragment of other serotypes.^19,21^ In addition, exchanging the binding domain of BoNT/X for H_C_/A produced a functional and efficient toxin.^8^ Altogether, the low toxicity of BoNT/X is likely linked to the unusual properties of its binding domain, including the hydrophobic surface and unclear receptor-recognition strategy, particularly towards gangliosides. This suggests an absence of suitable BoNT/X receptors on motor neurons of mammalian species.

BoNT/X was identified from a clinical isolate, but is evolutionary closer to other recently identified botulinum-like toxins, targeting insects (PMP1), or isolated from bovine feces (BoNT/En). It would therefore not be surprising if the true target of BoNT/X are non-mammalian species or non-neuronal cells. However, the medical and biotechnological potential of the functional LH_N_ fragment of BoNT/X is evidently vast and warrants further studies.^7,8,65^

## Materials and methods

### Purification of full-length, active BoNT/X

pK8-BoNT/X(Cloop)-10HT encodes BoNT/X full-length (UniprotKB P0DPK1) with the activation loop (C423 – C467) substituted with the activation loop of BoNT/C (Uniprot KB P18640, C437 – C453) and a C-terminus PreScission-cleavable H10 tag in the pK8 expression plasmid (pET26b derived synthesised at Entelechon). cDNAs were codon-optimized for *E. coli* expression and synthesized by GeneArt in 2 fragments for biosafety reasons: the light chain of BoNT/X (M1 – I422) with the activation loop of BoNT/C cloned into pK8 and the heavy chain of BoNT/X (I468 – D1306) with a C-terminal PreScission-cleavable H10 tag. The heavy chain was then cloned in pK8 in the C-terminus of the 1st fragment using BsaI for scarless insertion. NTNH/X (NCBI RefSeq WP_045538950) was cloned from pET28a-NTNH/X into pET32a using NdeI and XhoI restriction enzymes.

pET32a-NTNH/X and pK8-BoNT/X(Cloop)-10HT were co-transformed into NiCo21(DE3) competent *E. coli* (NEB). Protein expression was induced with 1 mM IPTG at 16 °C for 20 hours in mTB (Casein Digest Peptone 12 g/l, Yeast Extract 24 g/l, Dipotassium Phosphate 9.4 g/l, Monopotassium Phosphate 2.2 g/l, 0.4% Glycerol) supplemented with Kanamycin and Ampicillin. Cells were harvested at 4 °C and lysed by sonication in 50 mM HEPES pH 5.5, 300 mM NaCl, 10 mM Imidazole, treated with Benzonase nuclease (Sigma) and clarified by centrifugation. Proteins were purified using the ÄKTA Pure FPLC system (Cytiva). Cleared lysates were loaded into HisTrap HP columns (Cytiva). The protein complex was eluted with 50 mM HEPES pH 5.5, 300 mM NaCl, 500 mM Imidazole and separated by desalting into a higher pH buffer (50 mM HEPES pH 7.5, 300 mM NaCl) using a HiPrep Sephadex G-25 desalting column (Cytiva). After concentration to 1 mg/ml, activation was performed overnight at 4 °C with 5 μg of Factor Xa (NEB) per mg of protein. To separate activated BoNT/X(Cloop)-10HT from NTNH/X, the activated sample was loaded onto a HisTrap HP column equilibrated in 50 mM HEPES pH 7.5, 300 mM NaCl, and eluted in 2 steps at 500 and 1000 mM Imidazole. NTNH/X was found in the flow through and BoNT/X(Cloop)-10HT in elution fractions, which were pooled and buffer exchanged and adjusted to 0.1 mg/ml into PBS (Gibco) supplemented with 1 mg/ml BSA.

### Measurement of VAMP cleavage in cortical neurons

CTX cells were treated with serial dilutions of BoNT and incubated at 37 °C for 24 h, following 3 weeks *in vitro*. Cells were lysed by removing all medium and adding sample buffer (25% NuPAGE buffer (Life Technologies, Fisher Scientific) supplemented with 10 mM dithiothreitol and 250 units/μl Benzonase (Sigma)). Lysate proteins were separated by SDS-PAGE and transferred to nitrocellulose membranes. Primary antibodies used were against VAMP-2 (custom made) and VAMP4 (Santa Cruz sc-365332). The secondary antibodies were HRP-conjugated ant-rabbit IgG (Sigma A6154) and HRP-conjugated anti-mouse IgG (Sigma A4416). Proteins were visualised using an enhanced chemiluminescent detection system (Fisher Scientific). Luminescence detection was carried out using a Syngene GeneGnome and image analysis was performed using GeneTools software (Syngene Bioimaging, Cambridge UK). VAMP cleavage was monitored by measuring the disappearance of the specific full-length VAMP protein and the appearance of the cleaved fragment of the VAMP protein. The amount of cleaved VAMP protein was expressed as a percentage of the sum of full-length protein and cleaved product when available.

### Mouse Phrenic Nerve Hemi-Diaphragm assay

The mouse Phrenic Nerve Hemi-Diaphragm (mPNHD) assay is an isolated, *ex vivo* model used to measure the effect of botulinum neurotoxin (BoNT) at its *in vivo* target, the neuromuscular junction (NMJ). Male CD1 mice, 25-30 g (Charles River Laboratories, UK), were killed by CO_2_ asphyxiation and the hemi-diaphragm muscle and attached phrenic nerve isolated and attached to a custom tissue holder/electrode (Emka Technologies, France) installed in a 10 ml tissue bath (EmkaBATH4 Tissue Bath System, Emka Technologies, France) containing Krebs-Henseleit buffer (KHB; 118 mM NaCl, 1.2 mM MgSO_4_, 11 mM Glucose, 4.7 mM KCl, 1.2 mM KH_2_PO_4_, 2.5 mM CaCl_2_, 25 mM NaHCO_3_, pH 7.5 (Sigma, UK) at 37 °C) gassed with Carbogen (95% O_2_/5% CO_2_; BOC, UK). The phrenic nerve was continuously electrostimulated using 10 V, 1 Hz, 0.2 ms stimulation and resultant muscle contractions, as a result of Acetylcholine (ACh) release at the NMJ, were recorded using an isometric force transducer (Emka Technologies). The preparations were allowed to equilibrate for 45 mins in KHB renewed every 15 minutes. Following equilibration, the tissue was incubated with 3 μM tubocurarine hydrochloride (Sigma), a reversible, competitive antagonist at the nicotinic ACh receptor on the muscle cells. Subsequent inhibition of muscle contraction was considered an indication that the contraction response was due to ACh released by nerve stimulation (data from preparations with inhibition below 95% were discounted). The preparations were then washed 3× for 5 mins and after a further stabilization period of 20 mins, 1 ml of 10× BoNT (final concentration 10 pM for BoNT/A and BoNT/B (both LIST Biological Laboratories, Campbell, USA), and recombinant (r)BoNT/X, and 100 pM for LH_N_/X, in KHB supplemented with 0.05% (w/v) gelatin type A (Sigma) was added to the tissue bath and electrical stimulation continued until muscle contraction was ablated. Following complete paralysis, the muscle was directly stimulated (bypassing the phrenic nerve) by increasing the strength of the applied electro-stimulation (10 V, 1 Hz, 2 ms) in order to confirm the continued physiological viability of the preparation. Experimental data were recorded with IOX software (Emka Technologies). The decrease in contraction with time following toxin addition was calculated as a percentage of the contraction just before toxin addition and a four-parameter logistic curve fitted to the data, where appropriate, using GraphPad Prism (GraphPad Prism version 6.07 for Windows, GraphPad Software, La Jolla, CA, USA, www.graphpad.com). From the curve fitted to the data, the time to 50% diaphragm paralysis (t50) was estimated.

### Digital Abduction Score (DAS) Assay

All procedures were conducted in accordance with the guidelines approved by the Institute Animal Care and Use Committee (IACUC) at Boston Children’s Hospital. Female mice were purchased from Envigo (CD-1 strain, 17-20g). BoNTs were prepared in 0.2% gelatin-phosphate buffer (pH 6.3). Mice were anesthetized with isoflurane (3–4%) and injected with BoNT/A (6 pg) or BoNT/X (1 μg) by intramuscular injection into the gastrocnemius muscle of the right hind limb using a 30-gauge needle attached to a Hamilton syringe. Muscle paralysis and the spread of hind paw in response to a startle stimulus were observed and recorded. The hind limb digit abduction reflex in the mouse was induced by grasping the animal lightly around the torso and lifting it swiftly into the air or by lifting it with the nose pointing downwards. Animals were prescreened for a normal digit abduction response before the experiment and those showing abnormal digit abduction responses or hind paw deformities were excluded from the study. Typically, the percentage of animals with abnormal digit abduction responses is less than 1%. The digit abduction response of each mouse was scored live using a five-point scale, from normal reflex/no inhibition (DAS 0) to full inhibition of the reflex (DAS 4).^23^ Mice were scored for digit abduction response 8 h following BoNT injection, twice a day during the first week post-dosing, and once daily for 25 days, following injection of BoNT/A. We also measured body weight daily immediately after assessment of the digit abduction.

### Production of BoNT/X-NTNH/X for structural studies

The BoNT/X-NTNH/X complex was recombinantly co-expressed from a pET-22b vector encoding His_6_-tagged R360A/Y363F inactive mutant of BoNT/X and from a pET-28a(+) vector encoding NTNH/X (GenScript). The expression was performed in *E. coli* BL21(DE3) in Terrific Broth using the LEX bioreactor system (Harbinger Biotech, Toronto, Canada) and it was induced with 1 mM IPTG at OD_600nm_ of 1. The cells were harvested after around 20 hours of post-induction cultivation at 18 °C and lysed by sonication in lysis buffer (50 mM MES, 500 mM NaCl, 25 mM imidazole, 5% glycerol, 1 mM DTT, pH 5.5 plus complete EDTA-free protease inhibitor cocktail (Roche, Basel, Switzerland)). The BoNT-X/NTNH complex was purified by gravity-flow affinity chromatography on Ni-NTA agarose resin (Macherey-Nagel, Düren, Germany) in lysis buffer, the wash buffer contained 70 mM imidazole, and the his-tagged complex was eluted in three steps by 100 mM, 250 mM, and 500 mM imidazole in the lysis buffer. Fractions containing the BoNT/X-NTNH/X complex were pooled and dialyzed against the size-exclusion chromatography buffer (50 mM MES, 500 mM NaCl, 5% glycerol, 1 mM DTT, pH 5.5), concentrated, and further purified by size-exclusion chromatography (SEC) on a Superdex200 16/60 column and ÄKTA Pure system (GE Healthcare, Uppsala, Sweden). The pure BoNT/X-NTNH/X complex was concentrated to 8.4 mg/ml, flash frozen, and stored at −80 °C.

### SEC pH stability experiments

Fractions from the previous preparatory SEC with BoNT/X-NTNH/X were pooled, concentrated and loaded on a Superdex 200 16/60 column (GE Healthcare, Uppsala, Sweden) a total of five times, with the column equilibrated at different pH values (pH 5.5, 6.5, 7.5, 8.5, and 9.5). Each load contained approx. 0.25 mg of the complex. The SEC running buffers are listed below.

Buffer composition for SEC pH stability experiments.

| **pH** | **Buffer composition** |
| --- | --- |
| 5.5 | 50 mM MES, 150 mM NaCl, 0.5 mM TCEP |
| 6.5 | 50 mM MES, 150 mM NaCl, 0.5 mM TCEP |
| 7.5 | 50 mM HEPES, 150 mM NaCl, 0.5 mM TCEP |
| 8.5 | 50 mM Tris, 150 mM NaCl, 0.5 mM TCEP |
| 9.5 | 50 mM CHES, 150 mM NaCl, 0.5 mM TCEP |

### pH-dependent trypsin proteolysis

Recombinantly purified NTNH/X, BoNT/X, and the BoNT/X-NTNH/X toxin complex were subjected to limited proteolysis with trypsin for 13 h at room temperature. The trypsin digestions were performed at four different pHs in buffers containing 50 mM MES (pH 5.0), sodium phosphate (pH 6.0) or Tris (7.5 or 8.0) and 150 mM NaCl. The trypsin:protein sample ratios (w/w) were 1:5 (pH 5.0), 1:10 (pH 5.0 or 6.0) or 1:20 (pH 7.5 or 8.0): trypsin concentrations were raised at low pH conditions as trypsin showed reduced proteolytic activity under lower pH conditions. As controls, each protein was prepared and incubated in parallel, but without trypsin. The digestion was stopped by boiling samples in reducing SDS-loading buffer for 5 min. All samples were subjected to SDS-PAGE, followed by Coomassie blue staining and destaining.

### EM sample preparation

The BoNT/X-NTNH/X sample was diluted with 50 mM MES, 150 mM NaCl pH 5.5 and blotted onto the grids using the Vitrobot blotting robot (FEI, Hillsboro, OR, USA) at 100% humidity and 22 °C, waiting for 15 s for the sample to equilibrate before blotting for 1.5 s. Grids were clipped and stored in liquid nitrogen until further analysis. For the first data set, Quantifoil R2/2 Cu 400 mesh holey carbon grids (Quantifoil Micro Tools, Jena, Germany) coated with a 0.4 mg/ml graphene oxide suspension (Sigma-Aldrich, St. Louis, MO, USA) were used and the BoNT/X-NTNH/X concentration was 0.1 mg/ml. For the second data set, Quantifoil R1.2/1.3 Cu 300 mesh holey carbon grids (Quantifoil Micro Tools, Jena, Germany) without a graphene oxide coating were used and the BoNT/X-NTNH/X concentration was 0.5 mg/ml.

### EM data acquisition and processing

Two data sets were acquired on Titan Krios microscopes (FEI, Hillsboro, OR, USA) equipped with K2 summit direct electron detectors (Gatan, Pleasaston, CA, USA) and operating at 300 kV, using identical data collection parameters. The first data set was collected at the Stockholm node of the Swedish National Cryo-EM facility (SciLifeLab, Stockholm, Sweden) and the second data set at the Umeå node (Umeå University, Umeå, Sweden). Movies were acquired at 130,000× nominal magnification with a pixel size of 1.05 Å/pixel, defocus range of −1.8 to −3.4 μm in 0.2 μm steps, and a total exposure time of 10 s over 40 frames; the total dose was 35.6 and 35.9 electron/Å^2^ for the first and second data set, respectively. Frames were aligned, averaged, and dose-weighted in *cryoSPARC*.^66,67^ CTF estimation and downstream processing was also carried out in *cryoSPARC*. In the first data set, 2,885 movies were recorded, from which 70 were rejected. A total of 591,151 particles were automatically picked in cryoSPARC, and after 2D classification 100,920 particles were selected for 3D refinement. In the second dataset, 1,123,595 particles were automatically picked from 2,523 recorded movies, and 351,194 particles were selected in 2D classification and used for 3D refinement. Finally, the particle sets from both datasets were combined for 2D classification and subsequent 3D refinement, which included 432,063 particles and yielded the final map at 3.12 Å resolution, calculated based on the gold-standard FSC of 0.143.^68^

### Model building

The BoNT/X-NTNH/X complex model was built into the map using a combination of automated docking of the individual domains of homologous BoNT/A-NTNH/A complex (PDB code 3v0a)^13^ using Phenix^69^ and manual building in Coot,^70^ together with real space refinement in Phenix. Refinement was also carried out in Gromacs using the cryo-EM map as a restraint.^71^ The final cryoEM map and model coordinates were deposited into the PDB and EMDB databases under the accession codes 8BYP and EMD-16330, respectively. The cryoEM statistics are listed in Supplementary table 4.

### Ganglioside binding assay

Purified gangliosides GD1a, GD1b, GT1b, and GM1 (Carbosynth, Compton, UK) were dissolved in DMSO and diluted in methanol to reach a final concentration of 2.5 μg/ml; 100 μl were applied to each well of 96-well PVC assay plates (Corning; Corning, NY). After solvent evaporation at 21 °C, the wells were washed with 200 μl PBS/0.1% (w/v) BSA. Nonspecific binding sites were blocked by incubation for 2.5 h at 4 °C in 200 μl of PBS/2% (w/v) BSA. Binding assays were performed in 100 μl PBS/0.1% (w/v) BSA per well for 1 h at 4 °C containing protein samples (in triplicate) at concentrations ranging 6 μM to 0.003 μM (in serial 2-fold dilution). Samples consisted of the His6-tagged R360A/Y363F inactive mutant of BoNT/X, and a His6-tagged H_C_/A control. Following incubation, wells were washed three times with PBS/0.1% (w/v) BSA and incubated with an HRP-conjugated anti-6xHis monoclonal antibody (1:2000, ThermoFisher) for 1 h at 4 °C. After three washing steps with PBS/0.1% (w/v) BSA, bound samples were detected using Ultra-TMB (100 μl/well, ThermoFisher) as the substrate. The reaction was stopped after 10 minutes by addition of 100 μl 0.2 M H_2_SO_4_, and the absorbance at 450 nm was measured using an Infinite M200PRO plate reader (Tecan, Männedorf, Switzerland). Data were analyzed with Prism7 (GraphPad Software).

## Abbreviations

BoNT: Botulinum neurotoxin
NTNH: Non-toxic non-hemagglutinin protein
HC: BoNT heavy chain
LC, nLC: BoNT light chain, NTNH analogous light chain
H_N_, nH_N_: BoNT translocation domain, NTNH analogous translocation domain
H_C_, nH_C_: BoNT receptor-binding domain, NTNH analogous receptor-binding domain
H_CC_: C-terminal subdomain of H_C_
H_CN_: N-terminal subdomain of H_C_
LH_N_: BoNT light chain and translocation domain fragment
DAS: Digit Abduction Score assay
NAP: neurotoxin-associated protein
M-PTC, L-PTC: minimally functional progenitor toxin complex, large progenitor toxin complex

## Acknowledgements

We thank the Swedish National cryo-EM facility staff Marta Carroni, Julian Conrad, and Michael Hall for their help during the acquisition of the cryo-EM datasets.

## Funding sources

This work was supported by the Swedish Research Council (2022-03681, 2018-03406) and the Swedish Cancer Society (20 1287 PjF) (P.S.). This work was partially supported by Ipsen. The work of J.S. was supported from ERDF/ESF, OP RDE, project “IOCB Mobility” (No. CZ.02.2.69/0.0/0.0/16_027/0008477) granted to the Institute of Organic Chemistry and Biochemistry of the Czech Academy of Sciences. M.D. was partially supported by grants from National Institute of Health (NIH) (R01NS080833 and R01NS117626). M.D. holds the Investigator in the Pathogenesis of Infectious Disease award from the Burroughs Wellcome Fund. The data was collected at the Cryo-EM Swedish National Facility funded by the Knut and Alice Wallenberg, Family Erling Persson and Kempe Foundations, SciLifeLab, Stockholm University and Umeå University.

## Conflicts of interest

P.S. and M.D. are inventors on patents regarding BoNT/X. M.B., D.B., M.E., J.P. SD and F.H. are employees of Ipsen. The authors declare no additional conflict of interest.

