## Supplementary information for "Structure and activity of botulinum neurotoxin X"

### Supplementary Figure S1

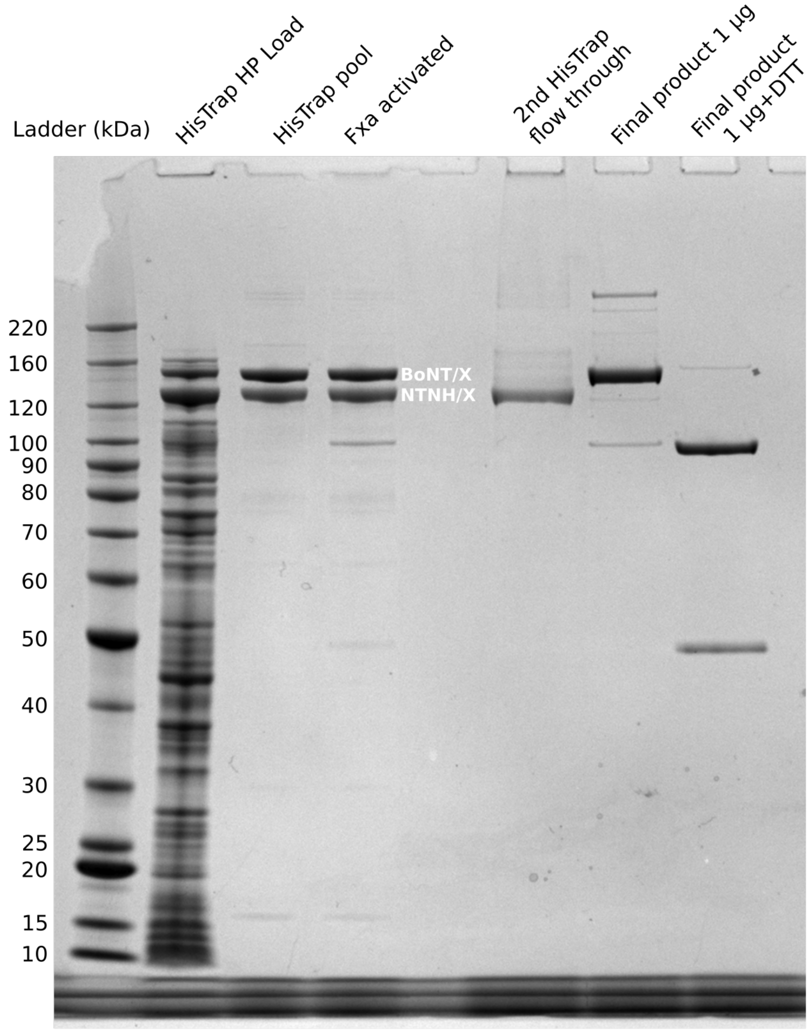

Supplementary Figure S1: Purification of BoNT/X. The lanes are marked at the top and the position of NTNH/X and BoNT/X are indicated in white. Molecular weight is indicated in kDa.

**Supplementary Figure S2**

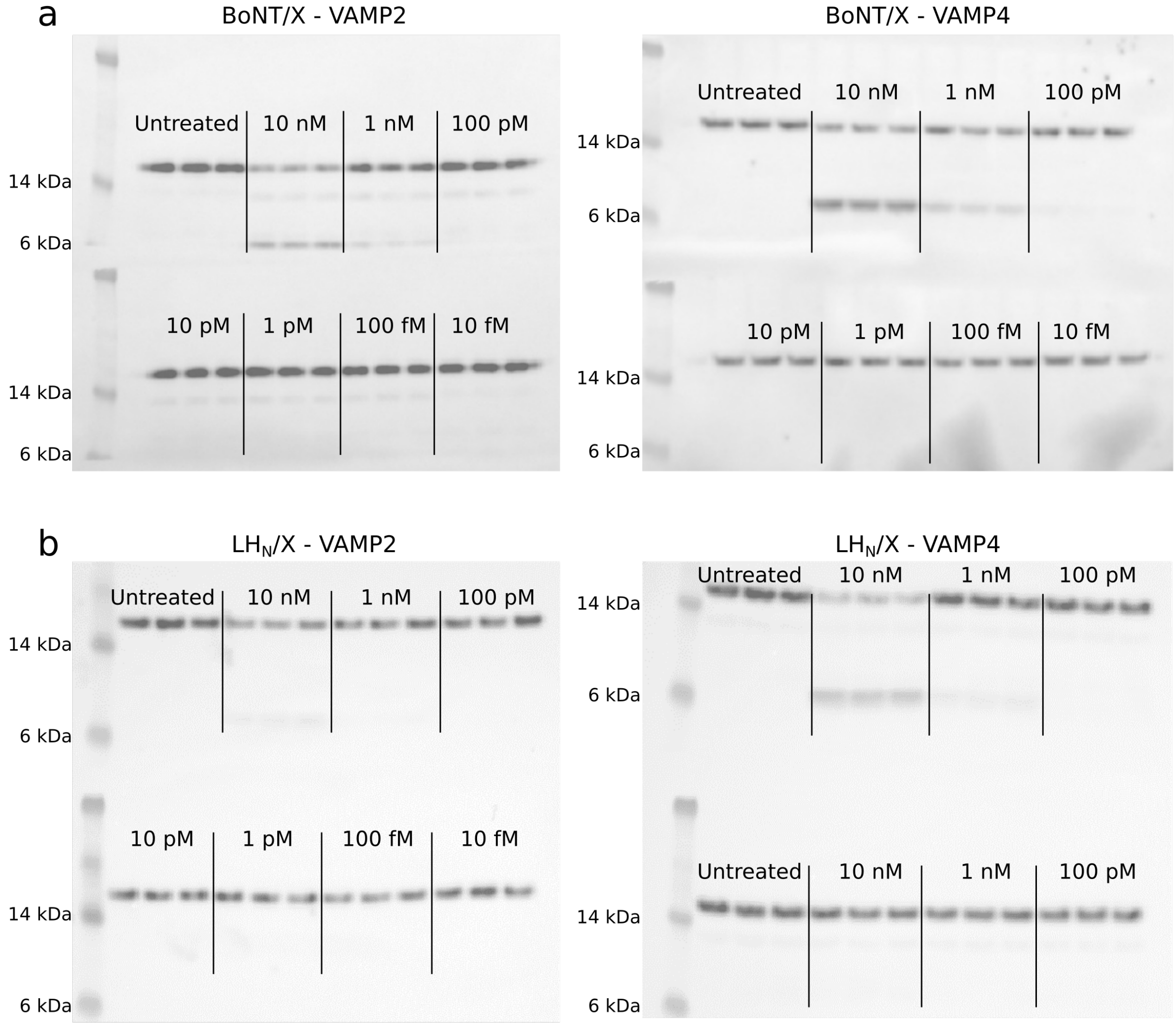

Supplementary Figure S2: Activity of BoNT/X and LH$_N$/X in cortical neurons. Rat cortical neurons were exposed to varying concentrations of recombinant full-length BoNT/X (A) or LH$_N$/X (B). After 24 h of incubation cells were lysed, and lysates analyzed for VAMP2 and VAMP4 by Western blots. VAMP migrates at about 15 kDa, whereas the VAMP cleavage product migrates at 7 kDa.

**Supplementary Figure S3**

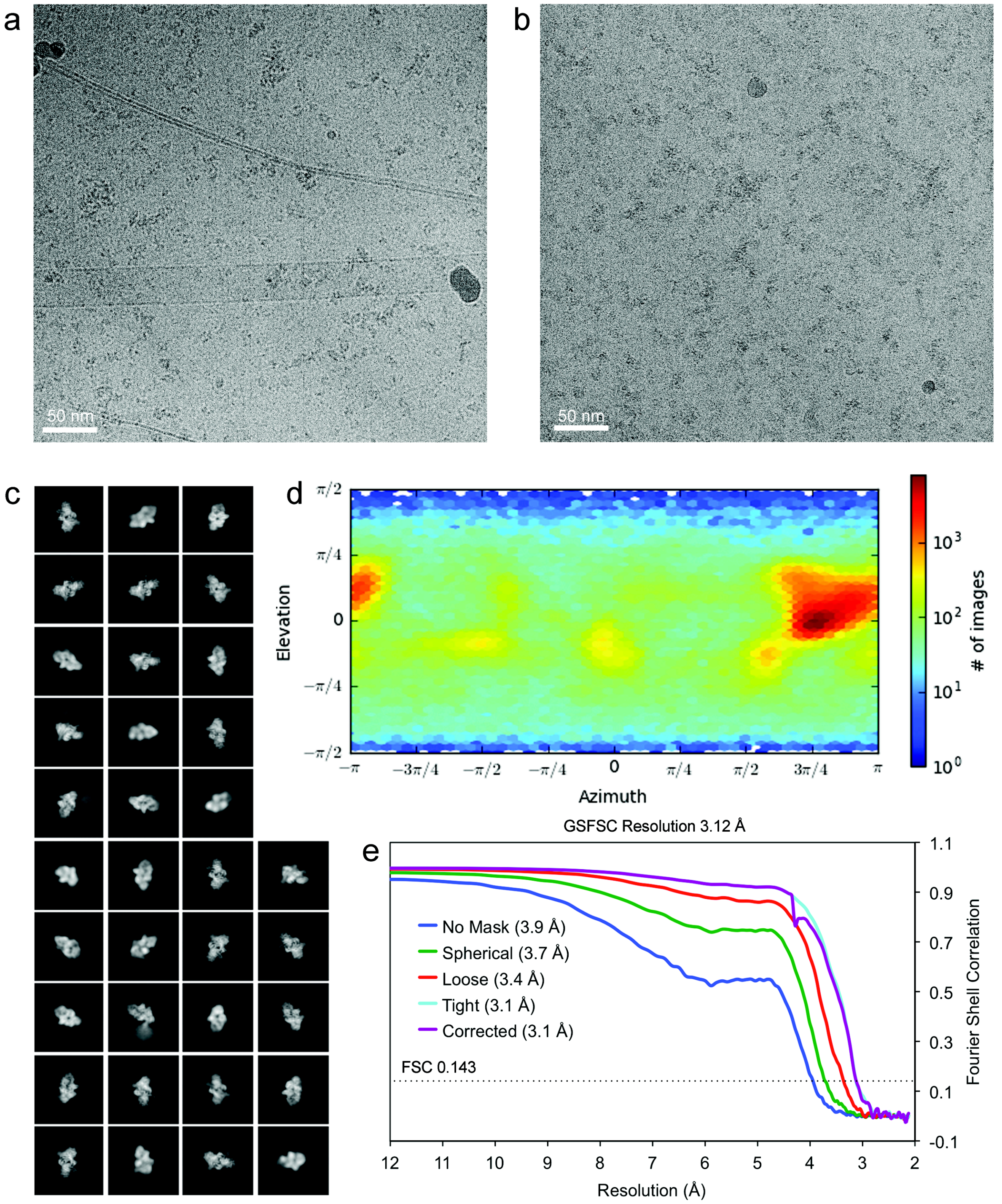

Supplementary Figure S3: cryoEM reconstruction of the BoNT/X-NTNH/X complex. Representative micrographs are shown from data set 1 (**a**) and data set 2 (**b**), together with final 2D classes (**c**). Panel (**d**) shows angular distribution of particle projections. The heat map shows number of particles for each viewing angle. (**e**) GSFSC plot for the final refined map (generated by cryoSPARC).

**Supplementary Figure S4**

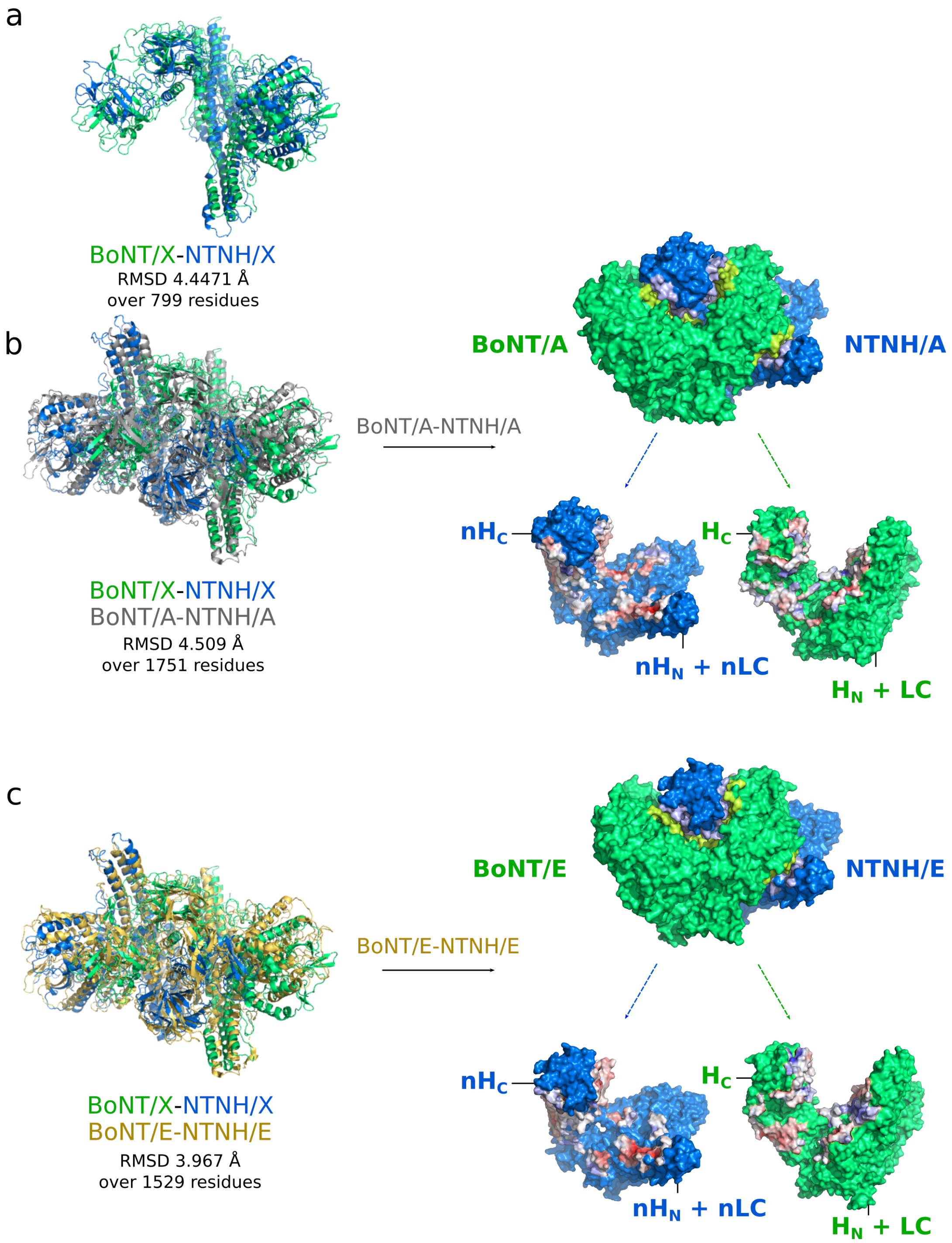

Supplementary Figure S4 (**a**) superposition of BoNT/X (green) and NTNH/X (blue), RMSD calculated over 810 Cα pairs. (**b**) superposition of BoNT/X-NTNH/X and BoNT/A-NTNH/A (PDB ID 3V0A, shown in grey). Acidic and basic residues on the interface are highlighted in red and blue respectively. (**c**) superposition of BoNT/X-NTNH/X and BoNT/E-NTNH/E (PDB ID 4ZKT, shown in yellow). Acidic and basic residues on the interface are highlighted in red and blue respectively.

### Supplementary Figure S5

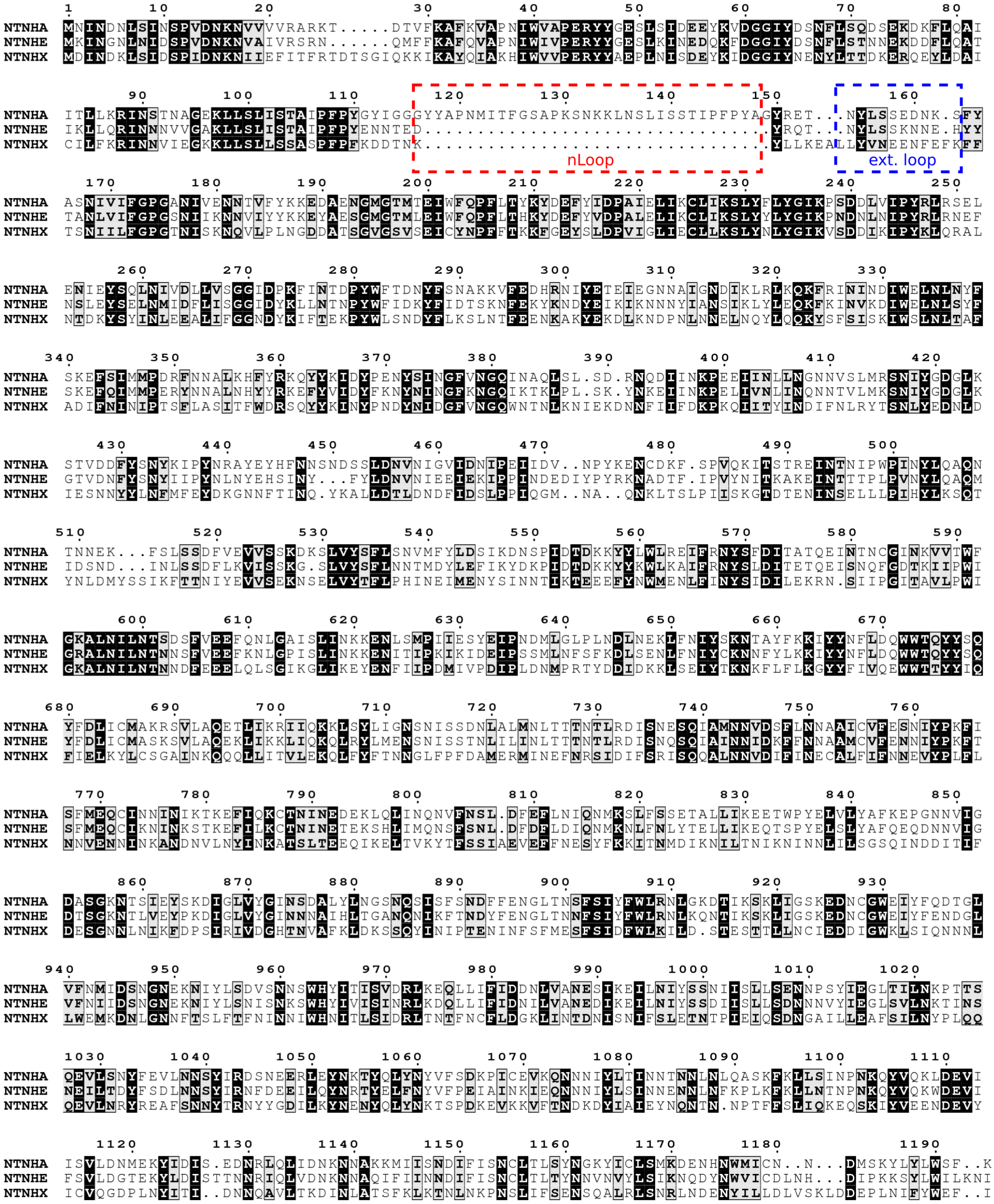

Supplementary Figure S5. Multiple sequence alignment of NTNH/A (UniProt Q45914), NTNH/E (UniProt P46082) and NTNH/X (NCBI RefSeq WP_045538950). The nLoop (residues 116 – 148) in NTNH/A is highlighted in a red box and the extended loop in NTNH/X with exposed hydrophobic residues, structurally close the NTNH/A nLoop (residues 128 – 139), is highlighted in a blue box.

**Supplementary table 1**

| BoNT/X dose | Mouse No. | Day 0 (DAS score/body weight) | Day 1 (DAS score/body weight) | Day 2 (DAS score/body weight) | Day 3 (DAS score/body weight) |
|---|---|---|---|---|---|
| 1 µg/mouse | #1 | 0/20 | 0/21 | 0/21 | 0/21 |
| | #2 | 0/20 | 0/22 | 0/22 | 0/21 |
| | #3 | 0/19 | 0/20 | 0/19 | 0/20 |

1 µg of BoNT/X (0.1 µg/µl) in 10 µl of PBS was injected into the mice leg using the DAS assay and the mice were monitored for 3 days. The mice did not show any paralysis.

**Supplementary table 2**

| BoNT/X dose | Mouse No. | Day 0 (DAS score/body weight) | Day 1 (DAS score/body weight) | Day 2 (DAS score/body weight) | Day 3 (DAS score/body weight) |
|---|---|---|---|---|---|
| 2 µg/mouse | #1 | 0/22 | 0/23 | 0/23 | 0/23 |
| | #2 | 0/29 | 0/29 | 0/29 | 0/29 |
| | #3 | 0/28 | 0/29 | 0/29 | 0/29 |

2 µg of BoNT/X (0.08 µg/µl) in 25 µl of 0.1 M phosphate buffer (pH 6.1) with 0.2% gelatin were injected into the mice leg using the DAS assay and the mice were monitored for 3 days. The mice did not show any paralysis.

**Supplementary table 3**

| | Mouse No. | Day 0 (body weight) | Day 1 (body weight) | Day 2 (body weight) | Day 3 (body weight) |
|---|---|---|---|---|---|
| BoNT/X | #1 | 23 | 23 | 23 | 24 |
| | #2 | 22 | 22 | 22 | 23 |
| | #3 | 22 | 21 | 22 | 22 |
| | #4 | 23 | 23 | 24 | 23 |
| Vehicle | #1 | 21 | 21 | 20 | 22 |
| | #2 | 22 | 23 | 23 | 24 |
| | #3 | 22 | 22 | 22 | 22 |

1 µg of BoNT/X was diluted in 100 µl of 0.1 M phosphate buffer (pH 6.1) with 0.2% gelatin and injected into CD-1 female mice intraperitoneally. A total of 4 mice were injected with BoNT/X and 3 mice were injected with vehicle only. The mice did not show obvious systemic symptoms or body weight loss.

**Supplementary table 4.** Cryo-EM data collection, refinement and validation statistics.

| | BoNT/X-NTNH/X (EMD-16330) (PDB 8BYP) | |
|---|---|---|
| **Data collection and processing** | | |
| Dataset | 1 | 2 |
| Magnification | 130,000× | 130,000× |
| Voltage (kV) | 300 | 300 |
| Electron exposure ($e^-/Å^2$) | 35.6 | 35.9 |
| Defocus range (μm) | -1.8 – -3.4 | -1.8 – -3.4 |
| Pixel size (Å) | 1.05 | 1.05 |
| Initial particle images (no.) | 591,151 | 1,123,595 |
| Final particle images (no.) | 432,063 | |
| Symmetry imposed | none | |
| Map resolution (Å) | 3.12 | |
| FSC threshold | 0.143 | |
| Map resolution range (Å) | 2.5 – 5.0 | |
| **Refinement** | | |
| Initial model used (PDB code) | 3v0a | |
| Model resolution (Å) | 2.9/3.3 | |
| FSC threshold | 0.143/0.5 | |
| Map sharpening *B* factor ($Å^2$) | -95.7 | |
| Model composition | | |
| Non-hydrogen atoms | 19,836 | |
| Protein residues | 2,421 | |
| *B* factors ($Å^2$) | | |
| Protein | 64.31 | |
| R.m.s. deviations | | |
| Bond lengths (Å) | 0.004 | |
| Bond angles (°) | 0.963 | |
| Validation | | |
| MolProbity score | 1.92 | |
| Clashscore | 9.76 | |
| Poor rotamers (%) | 0.31 | |
| Ramachandran plot | | |
| Favored (%) | 93.83 | |
| Allowed (%) | 6.17 | |
| Disallowed (%) | 0 | |

**Supplementary table 5**. Rmsd values for superpositions of $H_C$/X with other BoNT receptor-binding domains.

| **Structures** | **Rmsd (Å)** | **Cα pairs** |
|---|---|---|
| $H_C$/X – $H_C$/D (PDB ID 3OGG) | 1.653 | 345 |
| $H_C$/X – $H_C$/F (PDB ID 3FUQ) | 1.515 | 373 |
| $H_C$/X – $H_C$/E (PDB ID 4ZKT) | 1.980 | 356 |
